# Correlates of exercise hyperemia and muscle energetics in the human upper arm

**DOI:** 10.1101/2025.10.17.682308

**Authors:** J.A.L Jeneson, A.M. van den Berg-Faay, M.T. Hooijmans

## Abstract

We employed interleaved dynamic 1H magnetic resonance imaging (MRI) and 31P MR spectroscopy in combination with arm-cycling to investigate correlations of exercise hyperemia and muscle energetics in the triceps brachii (TB) muscle of the upper arm of healthy individuals. The parameter hyperemic slope (HS) determined from MRI acquisitions immediately after exercise was used as primary index of maximal TB oxygenation level in response to exercise. We found that HS tended to be inversely correlated with TB acidification during exercise (*P* =0.06) as opposed to findings in leg muscle. The absolute increase in cardiac-output during exercise was found to be uncorrelated with HS (*P* =0.19) suggesting that the magnitude of the hyperemic response to exercise involving a minor muscle mass is governed by local rather than systemic factors. Post-exercise, the rate of metabolic recovery was fastest in the study subject with highest HS and slowest for the opposite case, although this correlation failed to reach significance in our small study cohort (*P* =0.14). This finding fits the conclusion of previous musculoskeletal 31P MRS studies that oxygen supply to working skeletal muscle exerts significant metabolic control over oxidative muscular energy balance even if the physical task only involves a minor muscle mass.

**New & Noteworthy:** The use of interleaved dynamic 1H magnetic resonance imaging (MRI) and 31P MR spectroscopy uniquely afforded simultaneous interrogation of exercise hyperemia and muscle energetics in the human upper-arm. We found triceps muscle acidification during arm-cycling and readouts of hyperemia were, if anything, inversely correlated. Macrovascular and microvascular readouts of hyperemic response were uncorrelated. Post-exercise metabolic recovery rate tended to correlate with exercise hyperemia.

## Introduction

Perfusion of skeletal muscle can increase by one order of magnitude upon neural recruitment to support mechanical tasks such as exercise (1). The increase in muscular blood flow, more commonly termed hyperemia (1), is the result of a combination of coordinated central and local responses of the cardiovascular system including local vasodilation of feeding arterioles to balance the elevated muscular demand for oxygen towards mitochondrial ATP synthesis (1).

Various studies of muscle hyperemia in humans have documented substantial inter-individual differences in peak blood flow readouts in response to moderate intensity exercise tasks (2) or the surrogate exercise intervention of cuff inflation and deflation (3, 4). Yet, these reports have typically failed to raise any serious concerns about oxygen supply limitation of muscular metabolism impacting mechanical performance during submaximal exercise (e.g. (5)). Indeed, adequate oxygen availability to working muscle in healthy individuals has generally only been considered a concern during high-intensity isometric contractions (6) or during any high-intensity exercise involving a major muscle mass (7). However, Haseler et al observed in vivo that energy balance in calf muscle of healthy individuals performing voluntary single-leg plantar flexion at submaximal intensity improved almost instantaneously when subjects inspired oxygen-enriched air (8). Layec et al later found that amplification of the hyperemic response to exercise using cuff occlusion and release intervention accelerated post-exercise oxidative metabolic recovery in healthy subjects (9).

Here, we further investigated this subject matter in a cohort of healthy individuals using quantitative magnetic resonance imaging (MRI) and spectroscopy (MRS) techniques that afford simultaneous in vivo interrogation of muscle oxygenation level and oxidative ATP synthesis dynamics in response to concentric contractions involving only a minor muscle mass. Specifically, we employed 1H MRI to obtain quantitative readouts of the change in upper-arm muscle blood oxygenation in response to dynamic arm-cycling exercise (10), interleaved with dynamic in vivo 31P MRS readout of upper-arm muscle ATP metabolism and pH (11). To investigate the role of macrovascular changes in blood supply to hyperemia in working muscle (3) we additionally determined the increase in cardiac output in response to arm-cycling exercise in each subjects in a separate set of cardiac cineMRI measurements.

## Methods

### Study population

A total of six, normally active, healthy participants within the age range of 18 - 65 years were recruited for this study. All participants signed written informed consent.

### MR acquisition

MR datasets were acquired in the right upper arm on a 3T MR system (Ingenia, Philips Best, The Netherlands) using a previous described MR-compatible ergometer (10, 12) and a 6-cm diameter single loop ^31^P surface coil (Rapid Biomedical) fastened under the Triceps Brachii muscle. The MR examination consisted of a scout image to localize the arm and guide shimming and a time-resolved interleaved pulse acquire 31P MRS sequence (Adiabatic pulse; TR/TE: 1000/0.1ms; 2^nd^ order shimming) and multi-echo gradient echo acquisition (FFE; Axial slice 10mm; FOV 480x276; Acquisition matrix 160*92, 15 echoes; TR 27ms; TE/ΔTE 1.1/1.78ms; Flip Angle 15°) to simultaneous measure mitochondrial function and microvascular flow and oxygenation dynamics in rest, during in-magnet arm cycling and during subsequent recovery at a temporal resolution of 3.8 seconds and a total of 250 dynamics. Subjects were positioned supine head-first on the patient bed, the upper arm fixed to the bed, supported with sandbags and the elbow positioned in a 90° angle. After 30 seconds of rest, subjects were instructed to perform arm-cycling until exhaustion at 15W; thereafter recovery was mapped.

Additionally, we acquired cardiac triggered 2D CINE datasets (Velocity Encoded Phase Contrast (VE-PC) ; TR 4.48ms; TE 2.84ms; slice thickness 8 mm; acquisition matrix 140x136; pixel dimensions 1.21x1.21mm; Flip Angle 10; VENC 150 cm/s and progressively increased 40 frames) in rest and immediately after dynamic arm-cycling exercise to determine flow and volume differences in the ascending and descending aorta. The imaging plane is positioned orthogonal to the long axis of the ascending aorta at the level of the left ventricle.

### Data-processing and analysis

High-energy phosphate metabolism was assessed by quantitative evaluation of 31P MRS data. FIDs were analyzed using AMARES time domain fitting in jMRUI (13) using customized prior knowledge files (10). Level of Phosphorcreatine (PCr) depletion (%), end-exercise pH, and 95% Recovery Times (95%RT) of PCr (seconds) were calculated and used as metabolic outcome measures (10) (Figure 1 a & b).The post-exercise hyperemic response was characterized by analyzing the microvascular oxygenation dynamics using the offline reconstructed T2* maps of the upper arm muscles. The T2* maps were calculated from the multi-echo images using pixel-based mono-exponential fitting routine, excluding pixels with r-squared > 0.8 from the analysis (Figure 1d). Subsequently, T2* values were converted to R2* values (R2* = 1/T2*), which are often preferred for their more linear relationship with physiological parameters such as deoxyhemoglobin concentration. Regions of interest were manually delineated on the first echo image of the multi-echo gradient echo acquisition for the Triceps Brachii muscle using ITK-SNAP (14) (Figure 1c). The R2* values within each ROI were averaged and normalized to baseline (mean R2* over first 9 time points). Relative R2* values during rest, exercise and recovery were plotted over time. Post-exercise time-curves were analyzed using a customized composite function, consisting of a linear first part followed by an exponential second part. From the composite fit, three hyperemic outcome measures were derived including hyperemic peak (HP; %), the time-to-peak (TTP; sec) and the hyperemic slope (HS; -%/s)(Figure 1e). The hyperemic peak, reflecting a percentage change, was quantified according to:

**Figure 1.**
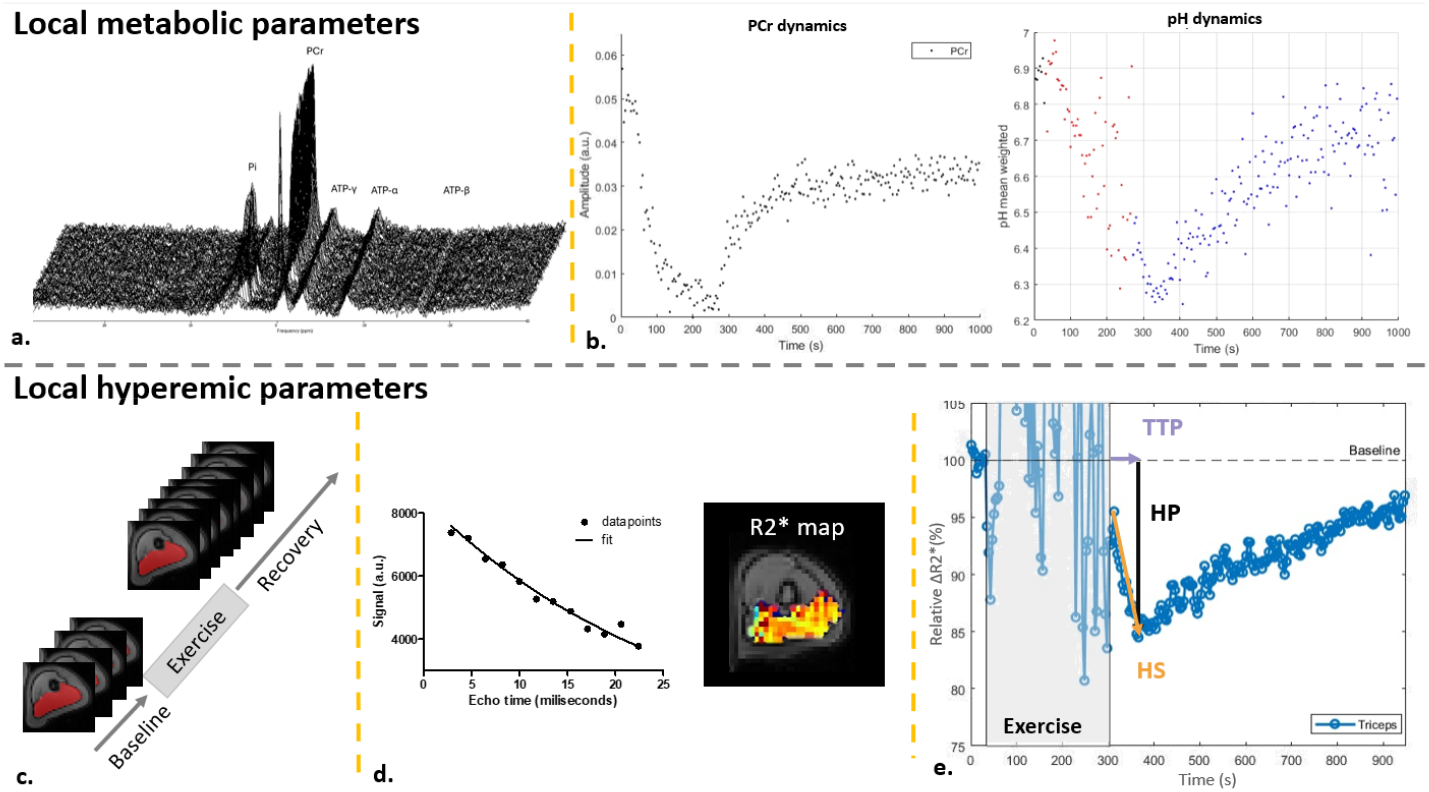
An overview of selected metabolic outcome parameters (top) and hyperemic outcome measures (bottom) is presented. (**a**.) A vertical stack of MR spectra from the triceps brachii muscle is shown for participant #4 at rest (t=0s), during arm-cycling exercise, and recovery (t=1000s). Key metabolite peaks are identified: HMP (Hexose-MonoPhosphates), Pi (inorganic Phosphate), PCr (PhosphorCreatine), and ATP (Adenosine TriPhosphate). (**b**.) Phosphocreatine and pH dynamics during rest, exercise, and recovery are visualized for participant #4. (**c**.) Gradient echo sequence images for this participant (bottom left) include manually segmented ROIs (red) for the triceps brachii muscle, used to characterize hyperemic outcome measures. (**d**.) A mono-exponential fit was applied to calculate the T2* map for the triceps brachii muscle. (**e**.) Mean normalized R2* values for the triceps brachii muscle are plotted over time to represent the hyperemic curves. Key outcome measures, including the hyperemic peak (HP) (black), Time-To-Peak (TTP) (Lila), and hyperemic slope (HS) (orange), are highlighted in the figure.

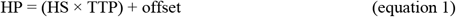

Central cardiovascular measures were derived from 2D VE-PC images acquired at rest and immediately following dynamic arm-cycling. These images were processed using Medis Suite 3.1 (Medis Medical Imaging Systems, Leiden, The Netherlands). Regions of Interest (ROIs) were manually delineated on the magnitude images across the time-series for the ascending and descending aorta. Forward flow volume and peak velocity were quantified based on these ROIs. Heart rate was recorded by the MR examiner during each phase of the protocol. The outcome measures included the change in heart rate (ΔHR) from rest to arm cycling, peak velocity and change in cardiac output volume between rest and arm cycling.

### Statistical Analysis

A combination of descriptive statistics and spearman correlation plots are used to describe the dataset. Significance level was set at 0.05.

## Results

### Participant characteristics and data-quality

All interleaved MR datasets, in 6 healthy participants (age range: 23-62 yrs. 3 male; normally active), were successfully acquired using this unique MR platform, resulting in good-quality metabolic time-curves and hyperemic response curves. Although all cardiac datasets were successfully acquired, one dataset could not be exported due to technical issues, resulting in 5 complete datasets. Additionally, the MR examiner reported delta heart rate for all participants in rest and directly after arm-cycling.

### Arm muscle hyperemic and metabolic measures and cardiac output measures

The range, mean, and standard deviation of individual outcome parameters are summarized in Table 1. The analysis of the metabolic time-curves and hyperemic response curves post-exercise revealed considerable variability between participants (Figure 2). A strong positive relationship was found between HS and TTP (r^2^=0.91; *P* =0.01). No significant relationship was found either between HP and TTP (r^2^=0.62; *P* =0.19) or HS and HP (r^2^=0.52; *P* =0.28) (Supplemental Figure 1). Substantial between subject variation was seen in the composition of the hyperemic peak signal. (Supplemental Table 1 and Supplemental Figure 2)

**Table 1.** Mean values, standard deviations (SD) and the range for all the local and central outcome measures used.

| Outcomes | | Range | Mean ± SD |
| --- | --- | --- | --- |
|  | Age (years) | 23 - 62 |  |
| **Metabolic** | PCr depletion (%) | 83 - 95 | 88.8 ± 5.3 |
| | End-exercise pH | 6.5 – 6.9 | 6.7±0.13 |
| | 95% PCr recovery times (seconds) | 76 - 548 | 216.4 ± 175.4 |
| | End-exercise pi/PCr | 4.1 - 14 | 8.2 ± 4.3 |
| **Hyperemic response** | Hyperemic Peak (%) | 14.8 - 34 | 24.1 ± 6.4 |
| | TTP (sec) | 32.5 - 119 | 65.3 ± 32.2 |
| | Hyperemic Slope (%/sec) | 0.05 – 0.27 | -0.16 ± 0.08 |
| **Systemic** | Delta HR (beats/min) | 5 - 35 | 17.8 ± 11.6 |
| | Volume difference (L/min) | 2.1 – 7.3 | 4.3 ± 2 |
| | Peak flow (cm/sec) | 103.8 – 199.7 | 149 ± 38.9 |

**Figure 2.**
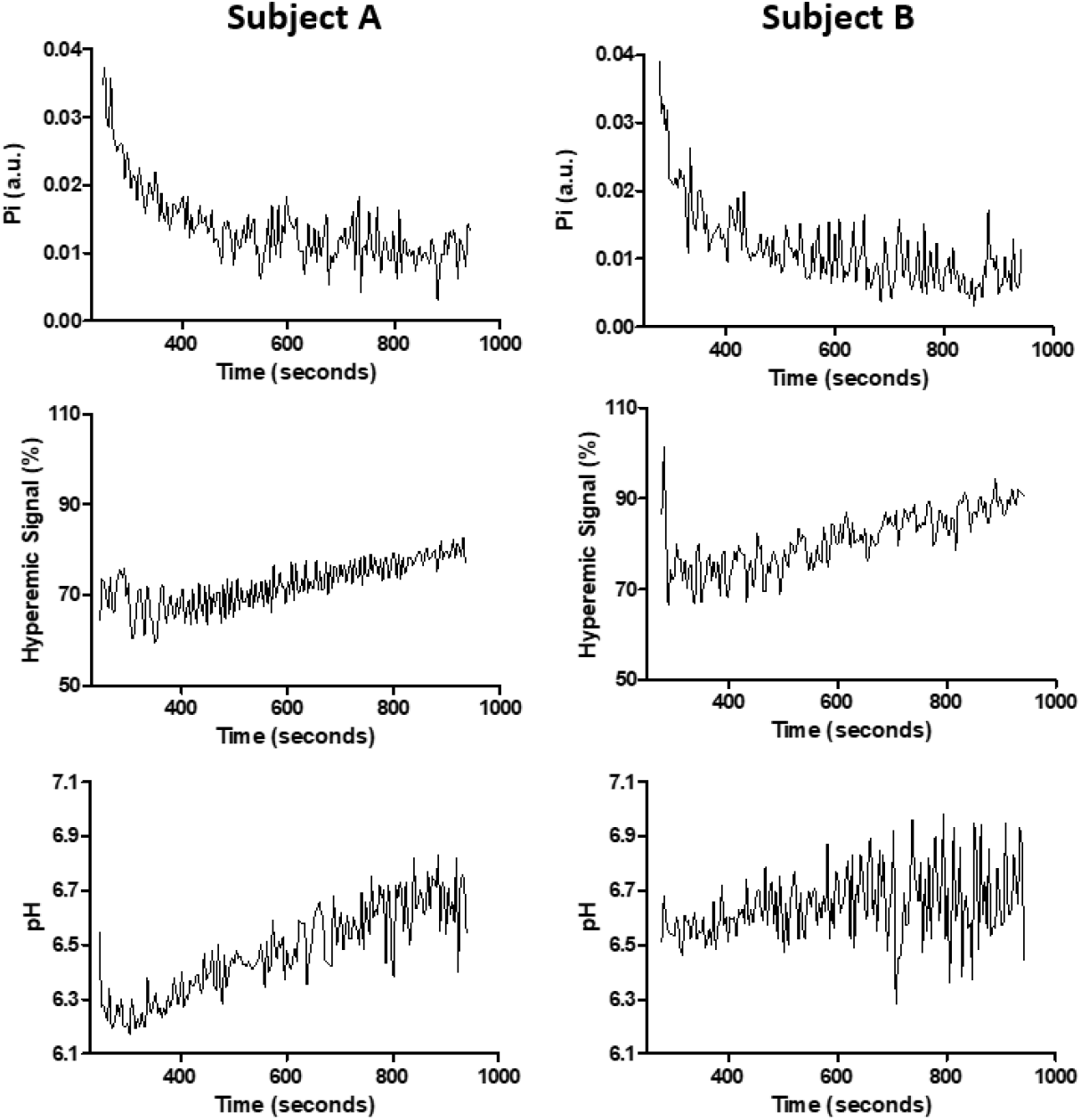
Time curves of Pi, normalized R2* signal and end-exercise pH post-exercise. Note the difference in patterns between two representative subjects for Pi, end-exercise pH and normalized R2* signal.

### Correlation Analysis of hyperemic measures, metabolic outcome measures and cardiac output measures in response to exercise

No significant correlations were found between hyperemic outcome measures and level of PCr depletion or end-exercise pH, although certain trends were noted (Figure 3). TTP and the HS demonstrated a negative association with end-exercise pH (TTP r^2^= -0.72; *P* = 0.11; HS r^2^= -0.8; *P* =0.06), suggesting that diminished local vascular responsiveness post-exercise may contribute to greater acidification. Conversely, no relationship was observed with the post-exercise hyperemic peak (r^2^=0.32; *P* =0.56), indicating that the magnitude of the hyperemic response may have a less critical role in this context. No clear correlations emerged between hyperemic response measures and cardiac output (HS r^2^= 0.70; *P* =0.19; TTP r^2^= 0.53; *P* =0.36; HP r^2^= -0.53; *P* =0.35)(Figure 4). Lastly, a positive trend was observed between 95% PCr recovery times and the HS (r^2^=0.67; *P* =0.14; Figure 5).

**Figure 3.**
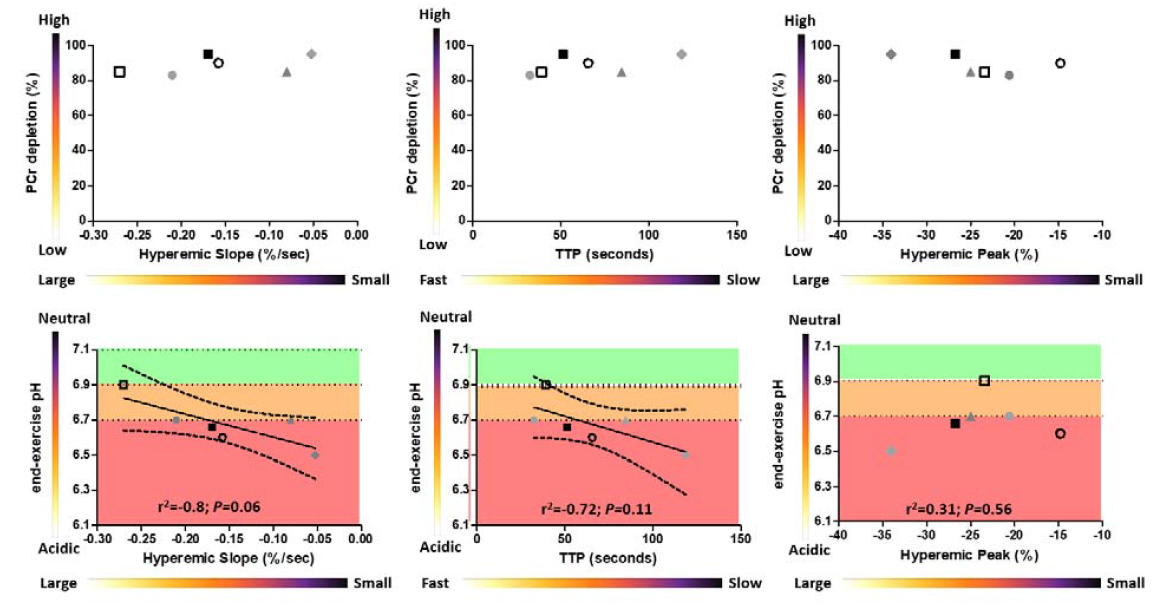
Scatter plots illustrate the correlation between PCr depletion (top row) and end-exercise pH (bottom row) with various hyperemic measures: hyperemic slope (HS) (left column), Time-To-Peak (TTP) (middle column), and hyperemic peak (HP) (right column). Each point represents data from an individual healthy participant, differentiated by unique symbols and/or colors. The pH ranges are visually delineated, with the neutral range highlighted in green, the intermediate range in orange, and the acidic range in red. Correlations are shown with a black line together with the 95% confidence intervals.

**Figure 4.**
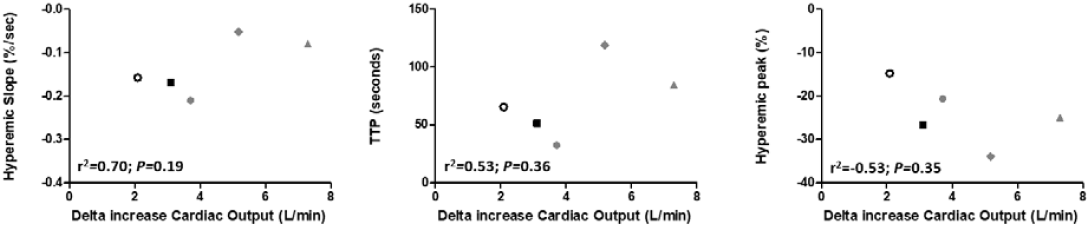
Scatter plots depict the correlation between cardiac output and key hyperemic measures: hyperemic slope (HS) (left column), Time-To-Peak (TTP) (middle column), and hyperemic peak (HP) (right column). Data points represent individual healthy participants, distinguished by unique symbols and/or colors.

**Figure 5.**
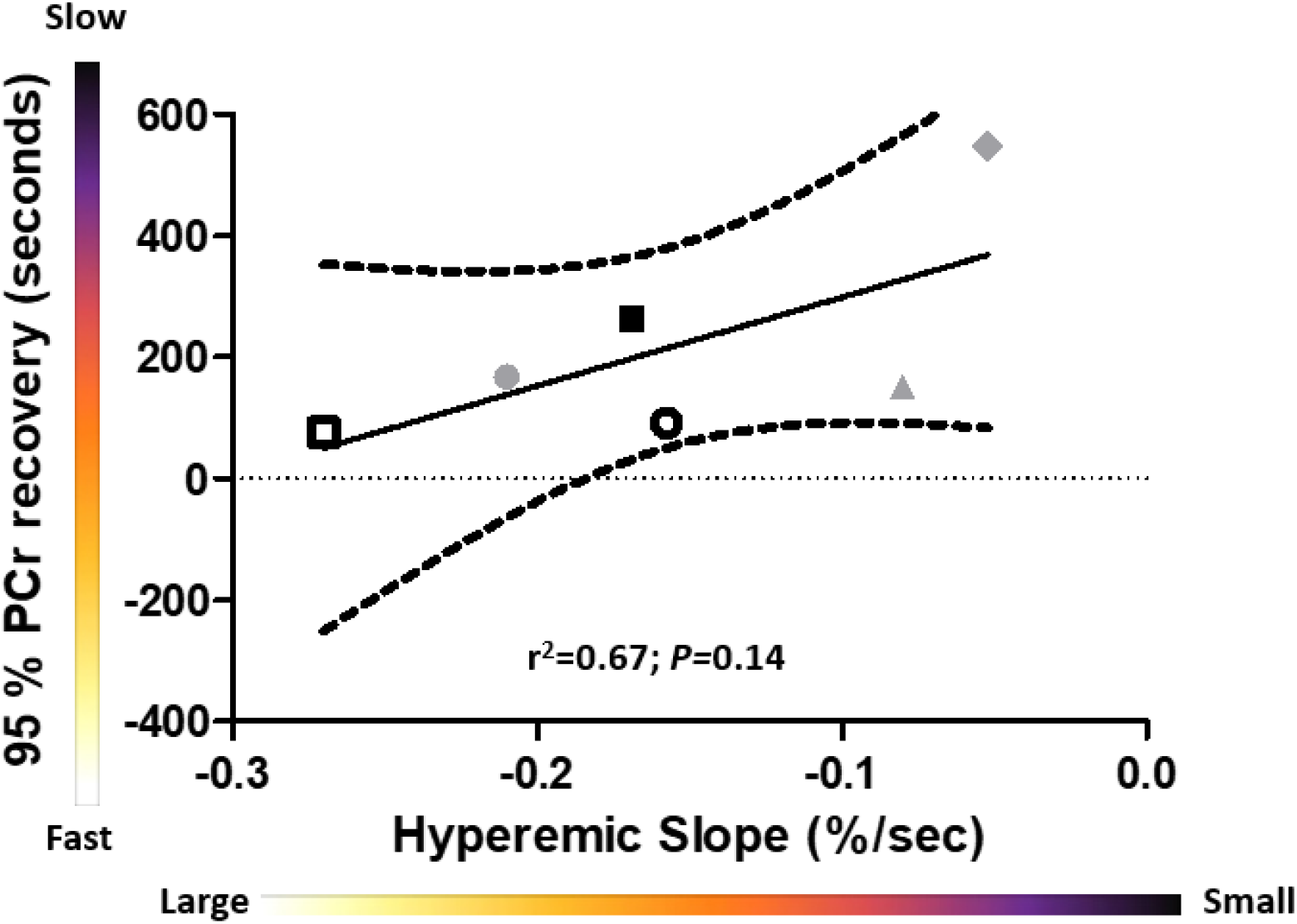
Scatter plot illustrating the correlation between 95% PCr recovery time and hyperemic slope (HS). Each point represents data from an individual healthy participant, differentiated by unique symbols and/or colors. Linear fit with 95% confidence intervals are shown with black lines in the graph.

## Discussion

The present study in a cohort of healthy individuals employed an interleaved 1H MRI/31P MRS sequence uniquely affording simultaneous tracking of both T2* as well as PCr, Pi and pH dynamics in the TB muscle of the upper arm in response to a bout of high-intensity arm-cycling. The former MRI parameter is an integrated magnetic resonance measure that reflects the net effect of blood volume, blood flow, and blood oxygenation levels on local magnetic field homogeneity (15, 16). The particular MRI pulse sequence that we used to measure T2* every ∼4s (see Methods) is, however, highly sensitive to motion artefacts (17). As such, we could not extract any quantitative information on TB blood oxygenation dynamics during the bout of arm-cycling exercise, including the key variable ‘peak blood flow’ in each subject. To overcome this problem, we exploited the fact that previous MRI recordings of muscle hyperemia have shown that there is typically a time lag on the order of tens of seconds after halting of exercise before muscle blood flow regresses to basal rate (18, 19). This afforded capture of the peak TB blood flow signature by qT2* MRI signal dynamics acquired from the moment exercise was halted and the arms were no longer moving. We indeed observed the post-exercise overshoot in hyperemic blood flow, oxygenation and volume manifesting as a continual drop in TB R2* for typically tens of seconds immediately after arm-cycling was halted (Figure 1e). We then fitted three parameters to capture the hyperemic overshoot in each subject (HS, HP and TTP, respectively; see Methods). Of these, we assumed that HS represented the best quantitative index of peak TB blood flow *during* exercise for two reasons. Firstly, we assumed that the particular fitted value of parameter TTP in an individual is, at least in part, also determined by the dynamics of restoration of basal vascular tone of the arterioles feeding the TB muscle that previously vasodilated during arm-cycling. Indeed, the particular value of HS in an individual was one and the same whether it was fitted over a time course equal to or shorter than TTP, respectively. However, we did find that HS and TTP were highly correlated (Supplemental Figure 1). Secondly, Figure 1e shows the value of parameter HP (in -%) is given by the equation HP = HS * TTP + offset. We assumed that variations in this offset term are primarily due to subtle differences in physical arm positioning within the static magnetic field pre-versus post arm-cycling, potentially influencing baseline signal intensity and local T2* (and/or T2) effects across subjects. Indeed, we found that latter parameter was uncorrelated with either HS or TTP (Supplemental Table 1).

Against this background, we recorded the fastest PCr recovery in the study subject with the highest value of HS (Figure 5). The inverse was also the case (Figure 5). In our small cohort of healthy individuals, the correlation of these parameters was moderately strong (r^2^=0.67) but failed to reach significance (*P*=0.14). Since post-exercise PCr recovery is principally driven by oxidative ATP synthesis (20), this outcome of our investigation provides independent support for the conclusion of two previous in vivo MRS studies from the Richardson lab that oxygen supply to working skeletal muscle constitutes an additional significant variable even when the physical task involves a minor muscle mass (8, 9). As such, they raise further doubts about, if not invalidate, the widely held view that in vivo 31P MRS recording of post-exercise PCr recovery kinetics in and by itself constitutes a robust quantitative assay of mitochondrial function in clinical investigations (5). Ideally, such examinations should include interrogation of muscle hyperemia dynamics in comparison to healthy controls using MRI or NIRS-based techniques.

Our combined MRI and MRS datasets from the TB muscle also afforded investigation of a previously reported close association between the post-exercise rate of change in human calf muscle pH and the signal intensity of an echo-planar MRI (EPI) sequence (21). The latter is likewise sensitive to changes in T2* but also T2 associated with changes in water content (17). Such a mechanistic link between muscle acidification and vascular tone of feeding arterioles in principle exists. A number of studies have indeed found that a change in pH of the interstitial space between working myofibers and the vascular bed has an inhibitory effect on upstream sympathetic vasoconstriction of their feeding arterioles in response to exercise (22-24). In the present study, however, we found that the time course of TB pH and HP immediately following exercise were uncorrelated (Figure 2). We did find a strong trend (*P=*0.06) towards significance for the correlation between end-exercise pH and HS (Figure 3). However, this particular correlation was negative instead of the expected positive correlation (22-24) – i.e., the largest absolute value of HS was found subjects with the lowest, not largest drop in TB pH during exercise. Within the constraints of our assumption that HS constitutes the best index of peak hyperemia during exercise, this result suggests that the contribution of sympatholysis to hyperemia in the TB muscle during arm-cycling was only minor compared to upstream reactive vasodilation of feeding arterioles. Indeed, Savard et al previously found that the increase in arterial and venous (nor)epinephrine levels during arm-cycling was only minor compared to leg-cycling (25).

Lastly, we investigated in our cohort of study subjects if any of the MRI measures of microvascular oxygenation dynamics in the TB muscle were correlated with macrovascular flow dynamics in response to the bout of arm-cycling exercise measured at the level of left-ventricular aortic outflow. Bopp and coworkers previously studied such a correlation between brachial arterial flow and forearm muscle hyperemia, respectively, and found that the kinetics, but not amplitudes of the increase in blood flow in the brachial artery and forearm muscle were closely correlated (3). We likewise failed to find any correlation between any of our 1H MRI indices of TB hyperemia and macrovascular blood flow during arm-cycling in our study cohort (Figure 4). This result suggested that the magnitude of the hyperemic response to exercise involving a small muscle mass such as arm-cycling is governed by local rather than systemic factors. This conclusion fits well with contemporary understanding of the principal controls of perfusive blood flow in working muscle (1).

This study has a number of limitations. Firstly, all data were collected at a relatively high workload (15-20 W; (12)) for the upper arm muscles of healthy subjects naïve to arm-cycling. This was caused by the fact that we employed an MR-compatible cycle ergometer with a relatively heavy wooden flywheel originally designed for leg-cycling (26) refitted for arm-cycling with a minimal workload on the order of 15 W (12). Secondly, our conclusions are based on a limited number of normally active healthy participants performing high intensity arm-cycling exercise. Therefore the generalizability of these findings to other populations or exercise intensities remains limited. Lastly, the signal intensity dynamics of our 1H MRI recordings from the TB muscle reflected the sum of temporal changes in blood flow, blood volume as well as oxygenation state in the interrogated cross-sectional slice rather than blood flow per se (17). Yet, a previous study interrogating muscle hyperemia likewise on basis of MRI data collected immediately following exercise found that the signal dynamics of an arterial spin labeling (ASL) MRI sequence and BOLD MRI were highly correlated (18). Since the former MRI technique exclusively measures blood flow (17) the findings of that study support our physiological interpretation of the 1H MRI results of the present study.

In summary, our investigation of hyperemic and metabolic dynamics in the TB muscle of the upper-arm in a small cohort of healthy individuals performing a bout of arm-cycling has found that the rate of oxidative metabolic recovery following exercise and our 1H MRI index of TB blood flow and oxygenation during exercise was moderately strong. That particular result supports previous findings (8, 9) that convective oxygen supply to working muscle is a determinant of its integral oxidative ATP synthetic capacity even in the presence of ample cardiac reserve. As such, our present findings reiterate the need for objectification of microvascular flow adequacy in clinical applications of post-exercise 31P MRS assay of PCr recovery kinetics to evaluate mitochondrial function in skeletal muscle.

## Supplemental figures

**Supplemental Figure 1.**
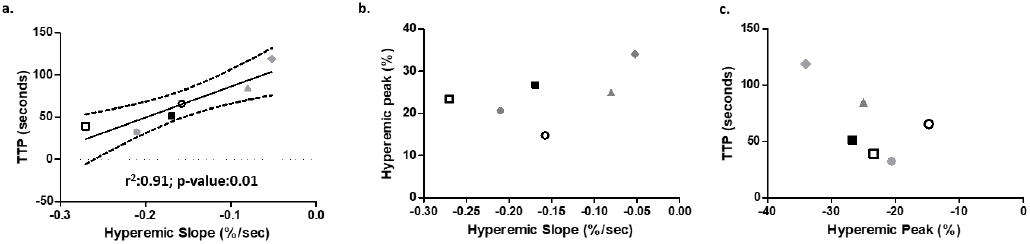
Scatter plots illustrate the correlation between Time-To-Peak (TTP) and hyperemic slope (HS) (left), hyperemic peak (HP) and hyperemic slope (HS) (middle) and hyperemic peak (HP) and Time-To-Peak (TTP) (right). Each point represents data from an individual healthy participant, differentiated by unique symbols and/or colors.

**Supplemental Figure 2.**
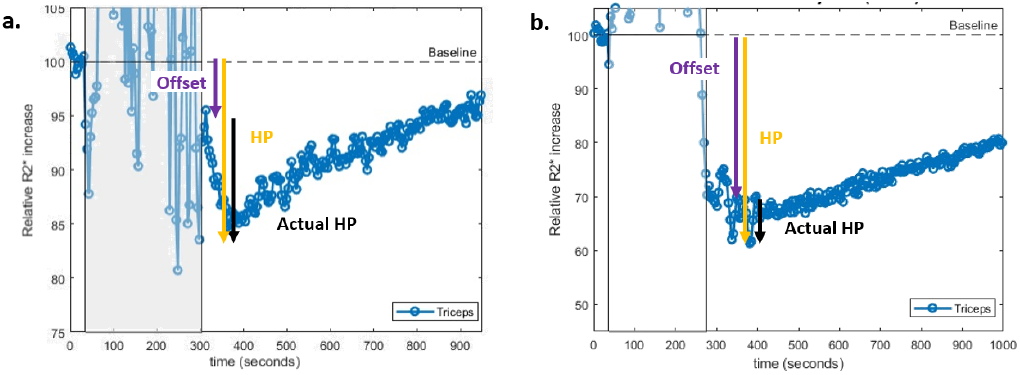
Mean normalized R2* values for the triceps brachii muscle are plotted over time to represent the hyperemic response curves for two subjects (**a-b**). The Hyperemic peak (HP) (orange), the offset (purple) and the actual hyperemic peak (black) are depicted in the figure.

**Supplemental Table 1.**

| Subject | HP (%) | Actual HP (%) | Offset |
| --- | --- | --- | --- |
| Subject 1 | -20.6 | -6.0 | -14.6 |
| Subject 2 | -26.7 | -8.7 | -18.0 |
| Subject 3 | -25.0 | -4.9 | -20.1 |
| Subject 4 | -34.0 | -4.0 | 30.0 |
| Subject 5 | -14.8 | -11.0 | 3.8 |
| Subject 6 | -24.4 | -9.4 | -15.0 |

Overview of the inter-subject variation in composition of the hyperemic peak (HP) signal.

## Acknowledgments

We would like to thank Aart Nederveen and Gustav Strijkers

## Author Contributions

Conceived and designed research: MTH, JALJ

Analyzed data: MTH, JALJ

Performed experiments: AMF, MTH, JALJ

Interpreted results of experiments: MTH, JALJ

Prepared figures: MTH, JALJ

Drafted manuscript: MTH, JALJ

Edited and revised the manuscript: AMF, MTH, JALJ

Approved final version of the manuscript: AMF, MTH, JALJ

