## Supplementary figures and images for "Correlates of exercise hyperemia and muscle energetics in the human upper arm"

### Supplemental Figure 1

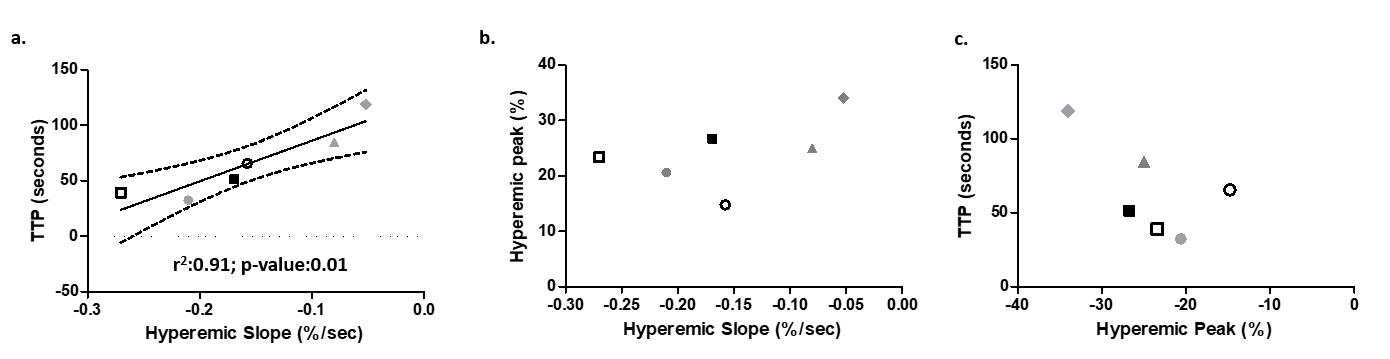

### Supplemental Figure 2

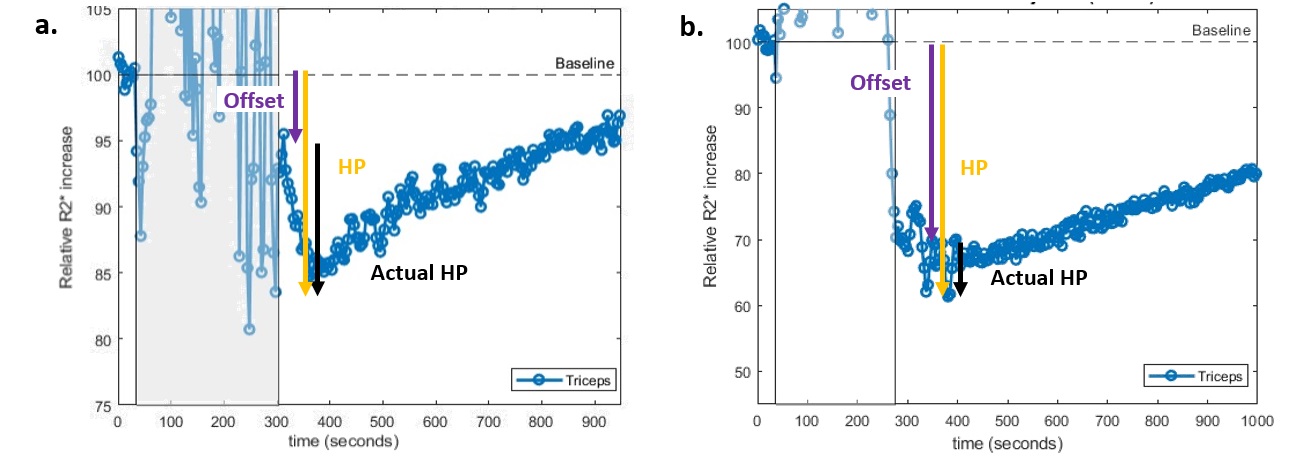
